# Dual receptive fields underlying target and wide-field motion sensitivity in looming sensitive descending neurons

**DOI:** 10.1101/2022.10.19.512946

**Authors:** Sarah Nicholas, Yuri Ogawa, Karin Nordström

## Abstract

Responding rapidly to visual stimuli is fundamental for many animals. For example, predatory birds and insects alike have amazing target detection abilities, with incredibly short neural and behavioral delays, enabling efficient prey capture. Similarly, looming objects need to be rapidly avoided to ensure immediate survival, as these could represent approaching predators. Male *Eristalis tenax* hoverflies are non-predatory, highly territorial insects, that perform high-speed pursuits of conspecifics and other territorial intruders. During the initial stages of the pursuit the retinal projection of the target is very small, but grows to a larger object before physical interaction. Supporting such behaviors, *E. tenax* and other insects have both target-tuned and loom-sensitive neurons in the optic lobes and the descending pathways. We here show that these visual stimuli are not necessarily encoded in parallel. Indeed, we describe a class of descending neurons that respond to small targets, to looming and to widefield stimuli. We show that these neurons have two distinct receptive fields where the dorsal receptive field is sensitive to the motion of small targets and the ventral receptive field responds to larger objects or widefield stimuli. Our data suggest that the two receptive fields have different pre-synaptic input, where the inputs are not linearly summed. This novel and unique arrangement could support different behaviors, including obstacle avoidance, flower landing, target pursuit or capture.

**Significance Statement:** If you are playing baseball, when the ball is far away, it appears as a very small object on your retina. However, as the ball gets closer, its image becomes a rapidly expanding object. Here, we show that within the hoverfly visual system, a single neuron could respond to both of these images. Indeed, we found a class of descending neurons with dual sensitivity, separated into two distinct parts of the visual field. The neurons have a more dorsal receptive field that is sensitive to small targets and a more ventral receptive field that is sensitive to larger objects.

## Introduction

The ability to respond quickly to visual stimuli is vital for the survival of many animals. Indeed, visual input may be used for a variety of tasks, from navigating around obstacles, choosing a suitable surface to rest upon, and also for detecting predators or prey. For example, many predatory insects rely on vision to identify suitable prey and engage in pursuits, doing so with astonishing precision and accuracy (e.g. Olberg et al., 2007; Nityananda et al., 2016; Fabian et al., 2018). Some non-predatory insects, including hoverflies, also have superb target detecting capabilities, which they may use for territorial defense or courtship (Fitzpatrick and Wellington, 1983; Zeil, 1986). Small target motion detector (STMD) neurons in the hoverfly optic lobe, and their presumed post-synaptic targets, the target selective descending neurons (TSDNs), have size and velocity tuning properties that match the target image at the start of the pursuit, suggesting that they could support these behaviors (Nordström et al., 2006; Nicholas et al., 2020; Thyselius et al., 2023).

In addition to the precise responses which occur during pursuit, behaviors elicited by looming stimuli, such as the escape response, also need to be fast and accurate (e.g. Fotowat et al., 2009; Santer et al., 2012; von Reyn et al., 2017; Mancienne et al., 2021; Lenzi et al., 2022). Indeed, neurons that respond strongly to rapidly looming stimuli exist in a range of species and visual structures, including the cat superior colliculus (Liu et al., 2011), the bullfrog optic tectum (Nakagawa and Hongjian, 2010), and in zebrafish retinal ganglion cells (Temizer et al., 2015). In insects, there are looming neurons in the optic lobes as well as in the descending pathways. Examples of these include the locust LGMD/DCMD system (Santer et al., 2012), the *Drosophila* Foma-1 neurons (de Vries and Clandinin, 2012), and the descending giant fiber (Fotowat et al., 2009).

Historically, insect looming neurons have been studied in the context of predator avoidance (e.g. Fotowat et al., 2009; Santer et al., 2012; von Reyn et al., 2017). However, there is emerging evidence that looming neurons also play a key role in pursuit behaviors. For example, silencing *Drosophila* Foma-1 neurons not only affects the escape response (de Vries and Clandinin, 2012), but also the ability of male flies to follow females during courtship (Coen et al., 2016). This is interesting as many looming neurons also respond strongly to small moving targets. For example, the locust LGMD/DCMD pathway was originally thought to play a role in object tracking (Rind and Simmons, 1992), and several looming sensitive neurons in the locust central complex also respond to small moving targets (Rosner and Homberg, 2013). Conversely, some dragonfly TSDNs respond not only to targets but also to looming stimuli (Frye and Olberg, 1995; Gonzalez-Bellido et al., 2013).

Taken together, this suggests that some neurons classically defined as either target or looming selective (Santer et al., 2008; Gonzalez-Bellido et al., 2013) respond to both. Like locusts (Santer et al., 2008) and dragonflies (Gonzalez-Bellido et al., 2013), hoverflies have descending neurons that respond to both looming stimuli and to small moving targets (Nicholas et al., 2020). We here investigate this dual sensitivity and find that these descending neurons have two distinct receptive fields, one more dorsal that responds selectively to the motion of small targets, and one more ventral that responds to larger objects, including sinusoidal gratings and high-contrast edges. We show that when the center of the ventral grating receptive field is to the right of the visual midline, the local motion sensitivity is to rightward motion, and vice versa. However, the preferred direction of the dorsal target receptive field and the ventral grating receptive field are not always the same. We also show that the two receptive fields receive separate input, from the pre-synaptic target pathway and the pre-synaptic widefield motion pathway, respectively, and when stimulated simultaneously the responses are not linearly summed. We hypothesize that the unique response characteristics of these neurons could be used in different behaviors.

## Materials and Methods

### Electrophysiology

We recorded from 98 looming sensitive descending neurons (Nicholas et al., 2020) in 94 male *Eristalis tenax* hoverflies, reared and maintained in-house as described previously (Nicholas et al., 2018a). At experimental time the hoverfly was immobilized ventral side up, using a beeswax and resin mixture, before an opening was made in the thoracic cavity. A small silver hook was used to elevate and support the cervical connective and a silver wire inside the opening served as a reference electrode.

Recordings were made from the cervical connective using a sharp polyimide-insulated tungsten microelectrode (2 MOhm, Microprobes, Gaithersburg, USA). Signals were amplified at 100x gain and filtered through a 10 – 3000 Hz bandwidth filter using a DAM50 differential amplifier (World Precision Instruments, Sarasota, USA), with 50 Hz noise removed with a HumBug (Quest Scientific, North Vancouver, Canada). The data were digitized via a Powerlab 4/30 and recorded at 40 kHz with LabChart 7 Pro software (ADInstruments, Sydney, Australia). Single units were discriminated by amplitude and half-width using Spike Histogram software (ADInstruments).

### Visual stimuli

Hoverflies were positioned perpendicular to and 6.5 cm away from the middle of a linearized Asus LCD screen (Asus, Taipei, Taiwan) with a mean illuminance of 200 Lux, a refresh rate of 165 Hz and a spatial resolution of 2,560 x 1,440 pixels (59.5 x 33.5 cm), giving a projected screen size of 155 x 138°. Visual stimuli were displayed using custom software written in Matlab (R2019b, Mathworks) using the Psychophysics toolbox (Brainard, 1997; Pelli, 1997). The stimuli were not perspective corrected. When values are given in degrees, this corresponds to the retinal size in the center of the visual field. Velocities are given in mm/s.

Potential looming sensitive descending neurons were initially identified based on their response to a small, black, moving target (left, Fig. 1A-D, and see Nicholas et al., 2020). Those neurons that also responded to a looming stimulus (Nicholas et al., 2020) were kept for further analysis. All of these neurons responded stronger to the looming stimulus than to an appearance control (Fig. 1E, F). The looming stimulus was a black circle on a white background, expanding over 1 s from 1° diameter to 118° (right, Fig. 1A, C), with a 10 ms rate of expansion (Fotowat and Gabbiani, 2007), also referred to as l/|v|. The appearance control was a black circle with 118° diameter that appeared and remained on the screen for 1 s.

**Figure 1.**
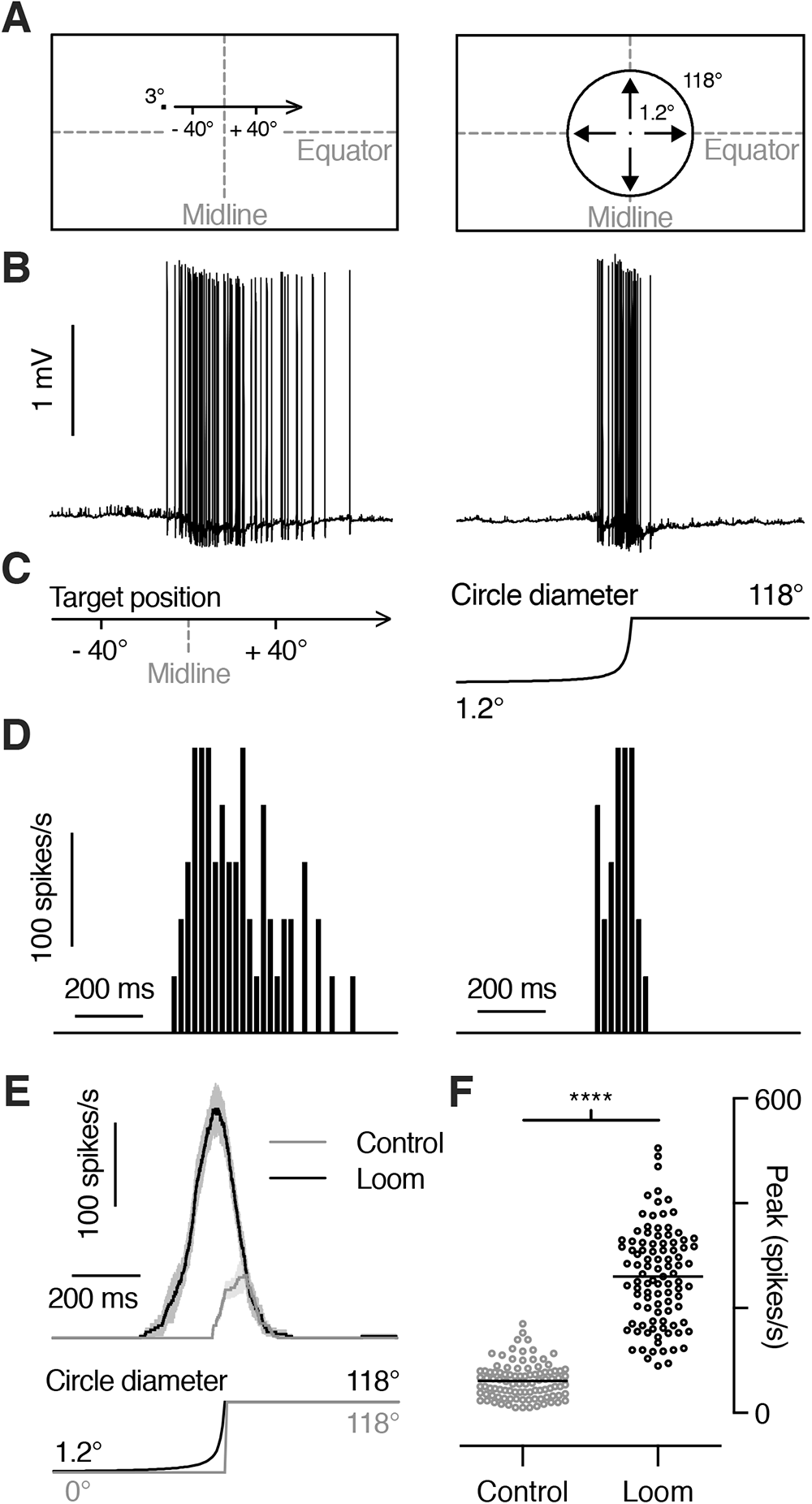
Looming sensitive descending neurons respond robustly to looming stimuli and to small targets. **A)** Pictograms of the 3° × 3° square target moving horizontally (*Left*) and the looming stimulus with an l/|v| of 10 ms and a final size of 118° (*Right*), as projected on the frontal visual monitor. **B)** Raw data trace from an extracellular recording of a looming sensitive descending neuron in response to a small target (*Left*) or a looming stimulus (*Right*). **C)** The position of the target on the visual monitor (*Left*) and the diameter of the looming stimulus (*Right*), time aligned with the data in panel B. **D)** Spike histograms of the responses in panel B using 20 ms bins. **E)** Example response from a single neuron (mean ± SEM, n = 4) to the looming stimulus and the appearance of a stationary black disc with a diameter of 118°. **F)** The peak amplitude of the response to a looming stimulus was significantly stronger than the peak response to the appearance control (p < 0.0001, paired t-test).

We mapped the target receptive field (Nicholas et al., 2020) of each neuron by scanning a target horizontally and vertically at 20 evenly spaced elevations and azimuths to create a 20 x 20 grid (Fig. S1A). The 3.5 x 3.5 mm (3 x 3°) black, square target moved at a velocity of 209 mm/s. As the screen width was larger than its height, the horizontal scans lasted for 2.8 s while the vertical scans for 1.6 s. There was a minimum 0.5 s interval between each stimulation.

We mapped the grating receptive field (Nicholas et al., 2020) using local sinusoidal gratings (93 x 93 mm, 71 x 71° in the visual field center, Fig. S1B) where the internal pattern moved in a series of 8 different directions presented in a pseudorandom order for 0.36 s each. The gratings had a wavelength of 17 mm (13° for the central patch, 0.08 cpd) and drifted at 5 Hz. The local gratings were placed in an overlapping tiling fashion so that 8 x 6 (width x height) squares covered the majority of the screen (Fig. S1B). There was a minimum 1 s interval between each stimulation.

To map the leading-edge receptive field, we scanned the entire height or width of the screen with an OFF-contrast edge moving left, right, down or up, at 209 mm/s (Fig. S3A-D).

For size tuning experiments a bar drifted at 209 mm/s, in the preferred direction of each neuron’s target receptive field. The bar drifted either horizontally or vertically through the center of the target or grating receptive field, as specified. The bar side parallel to the direction of travel was maintained at a fixed size of 3.5 mm (3°) whilst the perpendicular side varied from 0.2° to 155° (1 pixel to the full height of the screen). When presented simultaneously with a small target, the bar moved in the preferred horizontal direction through the grating receptive field only. In this case the bar height was varied between 5.7° and 106° (6.5 to 174 mm), whilst a fixed size 3 x 3° target moved through the target receptive field.

To determine the input mechanism of each receptive field, an OFF edge, an ON edge and a discrete bar, with a width of 3.5 mm (3°) drifted horizontally at 209 mm/s across the entire width of the screen. The height of these objects was 3° (3.5 mm) when drifted through the target receptive field, 84° (116 mm) when drifted through the grating receptive field, or the height of the screen (138°) to cover both receptive fields.

All stimuli were presented in a random order, except for the receptive field stimuli, which were presented in a pseudo-random order.

### Experimental Design and Statistical Analyses

All data analysis was performed in Matlab and GraphPad Prism 9.3.1 (GraphPad Software, USA). Statistical analysis was done using either GraphPad Prism 9.3.1 or the circular statistics toolbox (Berens, 2009) in Matlab, as appropriate. The sample size, statistical test and P-values are indicated in each figure legend, where *n* refers to the number of repetitions within one neuron and *N* to the number of neurons. Neurons were initially identified based on the response to a small target, with data from all neurons that subsequently passed the definition of a looming neuron (Nicholas et al., 2020, and see above) included in the analysis.

Even if the hoverfly was placed ventral side up in the experiments, all receptive fields are shown with the hoverfly’s dorsal side up. For target receptive field mapping we used the resulting 20 x 20 grid (Fig. S1A) to calculate the local preferred direction and local motion sensitivity (Fig. S1C), assuming a neural delay of 20 ms (Leibbrandt et al., 2021). We calculated the local average spike rate to the four directions of motion (dotted line, Fig. S1C) after subtracting the spontaneous rate, averaged in the 485 ms prior to stimulus presentation. We interpolated this to a 100 x 100 grid to generate receptive field maps (Fig. S1E) using Matlab’s *contour* function. We defined the center of the receptive field (Fig. S1E) as the center of the 50% contour line using Matlab’s *centroid* function. We fitted a cosine function (Nicholas et al., 2020) to the response to the four directions of motion (Nordström et al., 2006) and extracted its local preferred direction and amplitude (Fig. S1C). We calculated the preferred direction for each neuron by averaging the local preferred directions from the locations where the local motion sensitivity was above 50% of the maximum (blue, Fig. S1G).

For the grating receptive fields we used the resulting 8 x 6 grid (Fig. S1B) to quantify the local mean spike rate for each direction of motion, after removing the first 100 ms of the response, to avoid initial onset transients (Nordström and O’Carroll, 2009). We calculated the local mean response (dotted line, Fig. S1D) after subtracting the spontaneous spike rate, averaged during 800 ms preceding stimulus presentation. We spatially interpolated this 10 times and calculated the center from the 50% contour line (Fig. S1F). For each spatial location we fitted a cosine function (Fig. S1D) to the response to get the local preferred direction and local motion sensitivity (Nicholas et al., 2020). We calculated the overall direction selectivity using the top 50% of the local preferred directions (red, Fig. S1H).

We calculated the horizontal and vertical distance between the receptive field centers and the midline and equator. The distance between the two receptive field centers was calculated using the Euclidean distance.

For leading-edge receptive field mapping we first quantified the spike rate histogram for each neuron, after smoothing the spike rate with a 100 ms square-wave filter with 40 kHz resolution (Fig. S3A-D). We identified the maximum response to any direction of motion (purple, Fig. S3D). 50% maximum response was used as a threshold to determine the limits of the leading-edge receptive field (cyan, Fig. S3A, C, D). If a neuron’s response did not reach threshold to one direction of motion (e.g. Fig. S3B), the opposite direction of motion determined the receptive field outlines. If a neuron responded to both directions of motion (e.g. up and down) the outer thresholds were used to delineate the receptive field (Fig. S3E). From the resulting rectangular receptive field, we determined the center, and the proximity to the target and the grating receptive fields respectively (Fig. S3F), using the following proximity index:

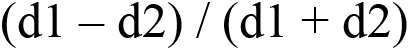

where *d1* is the Euclidean distance between the leading edge and the target receptive field centers, and *d2* the Euclidean distance between the leading edge and the grating receptive field centers (Fig. 4B). Thus, if the leading-edge receptive field center was closer to the grating receptive field center, the proximity index was positive, but if it was closer to the target receptive field center the proximity index was negative.

For all stimuli other than receptive field mapping, quantification of responses was done by averaging the spike rate within a 0.56 s analysis window centered on each neuron’s target or grating receptive field center, as specified.

### Code Accessibility

All Matlab scripts used for data analysis in this paper can be found here: https://doi.org/10.5281/zenodo.7227236

Private link for peer review: https://datadryad.org/stash/share/jZnw3f4ZNuytFZjEqDp2WNyAmKo32GGWTrWgTnywJaU

### Data Accessibility

All raw and analyzed data presented here have been deposited to DataDryad: https://doi.org/10.5061/dryad.6wwpzgn2p

Private link for peer review: https://datadryad.org/stash/share/jZnw3f4ZNuytFZjEqDp2WNyAmKo32GGWTrWgTnywJaU

## Results

### Looming neurons have dorsal target receptive fields and ventral grating receptive fields

We recorded from 98 looming sensitive descending neurons in male *Eristalis tenax* hoverflies. The neurons described here responded both to small target motion (left, Fig. 1A-D) and to looming stimuli (right, Fig. 1A-D). The response to a looming stimulus (right, Fig. 1B, D) started well before the stimulus reached its full size (right, Fig. 1C), and was much stronger than the response to an appearance control (Fig. 1E, F, and see Nicholas et al., 2020).

To investigate this dual sensitivity (Fig. 1A-D) in more detail, we mapped the receptive fields using two different methods. For this purpose we either scanned the visual monitor with a small, black target (Nordström et al., 2006) moving in four different directions (Fig. S1A, C), or we used a local sinusoidal grating (Fig. S1B, D) where the internal pattern drifted in eight different directions (Nicholas et al., 2020). The data from an example neuron show two distinct receptive fields, with a dorsal target receptive field (blue, Fig. 2A, S1E), and a ventral grating receptive field (red, Fig. 2A, S1F).

**Figure 2.**
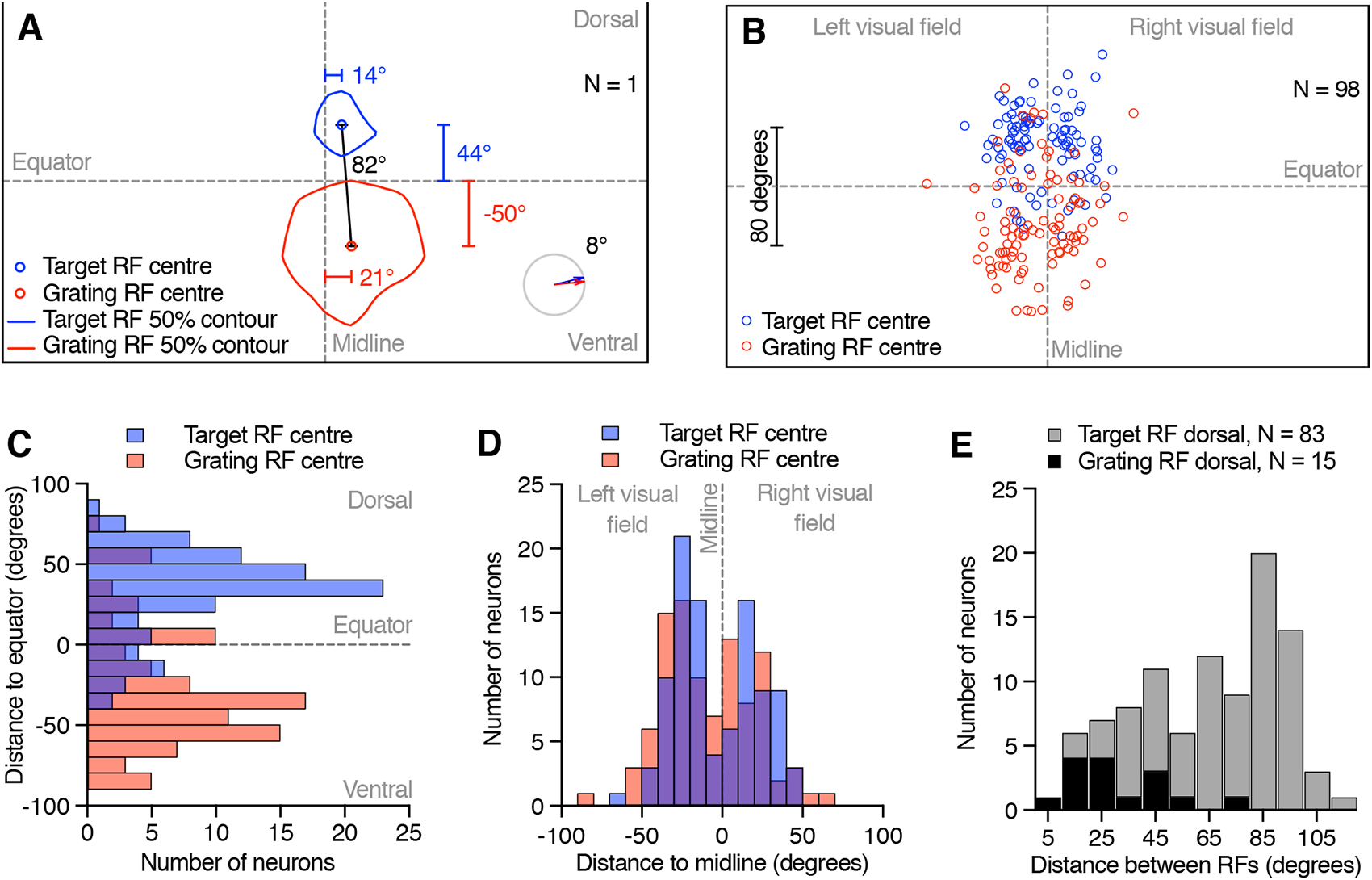
Two different receptive fields. **A)** The location of the two receptive fields of an example neuron as projected onto the visual monitor. The outlines show the 50% response and the small circle the center of the target receptive field (blue) and the grating receptive field (red). Euclidean distance between the receptive field centers (black line and value) and the distance of each receptive field center to the equator and the visual midline (colored lines and values) are indicated. Bottom right pictogram indicates the preferred direction of the target (blue arrow) and the grating receptive field (red arrow), and the difference between the two (black number). **B)** Location of target (blue) and grating receptive field centers (red) across 98 neurons. **C)** Vertical distance between target (blue) and grating receptive field centers (red) and the visual equator (10° bins). The target receptive field center locations were significantly different from the grating receptive field center locations (p < 0.0001, Mann-Whitney test). **D)** Horizontal distance from the visual midline (10° bins). There was no significant difference between the target receptive field center distance to the midline, and the grating receptive field distance (Mann-Whitney test). **E)** Euclidean distance between each neuron’s two receptive field centers (10° bins). Grey data come from neurons where the target receptive field was dorsal to the grating receptive field center (N = 83) and black data from neurons where the grating receptive field was dorsal to the target receptive field center (N = 15). When the grating receptive field center was dorsal (black), the distance between the two was significantly smaller (p < 0.0001, Mann-Whitney test).

We used the 50% response contours to locate the two receptive field centers (blue and red circle, Fig. 2A, S1E, F). Across the 98 neurons, the target receptive field centers cluster above the visual equator (blue, Fig. 2B), whereas the grating receptive field centers cluster below the equator (red, Fig. 2B), even if there are some exceptions (see also Fig. S2). We quantified the vertical distance between each receptive field center and the visual equator (44° and -50° in the example neuron, Fig. 2A), and found a bimodal distribution with target receptive field centers peaking 36° dorsal (median value) and grating receptive field centers peaking -36° (ventral, Fig. 2B, C).

We next quantified the horizontal distance between each receptive field center and the visual midline (14° and 21° in the example neuron, Fig. 2A). Across neurons we found a bimodal distribution with a gap along the visual midline (Fig. 2B, D). The grating receptive field center medians were at -27° and 19°, whereas the target receptive field center medians were at -25° and 20° (Fig. 2B, D). There was no significant difference between the target and the grating receptive field center distributions (Mann-Whitney test, left visual field, P = 0.31, right visual field, P = 0.40).

We noted that the target receptive field was often in the dorsal visual field, but that there were some exceptions (Fig. 2B, C, S2). We next investigated if the target receptive field was always dorsal to the grating receptive field and determined the Euclidean distance between the two receptive field centers (black line, 82°, Fig. 2A). Across neurons we found that the target receptive field was indeed most often dorsal to the grating receptive field (grey, Fig. 2E), and that the median distance between the two was 77°. When the grating receptive field was more dorsal, the two receptive field centers were significantly closer to each other (black data, Fig. 2E, median distance 26°, P < 0.0001, Mann-Whitney test).

### The grating receptive field is sensitive to motion away from the midline

We next determined the local motion sensitivity and average preferred direction of each neuron’s target and grating receptive field (colored arrows, Fig. S1G, H, S2). In the example neuron, the preferred direction of the target receptive field is toward the right (blue arrows, Fig. 2A, S1G), similar to the preferred direction of the grating receptive field (red arrows, Fig. 2A, S1H). For comparison across neurons we color coded the preferred direction into four cardinal directions, and plotted them as a function of receptive field center location. This analysis shows that the preferred direction of the target receptive fields depended on location (Fig. 3A). We found a significantly non-uniform distribution (P < 0.01, Rayleigh test), with a median direction preference up and away from the visual midline (vector lengths 0.37 and 0.37, insets, Fig. 3C).

**Figure 3.**
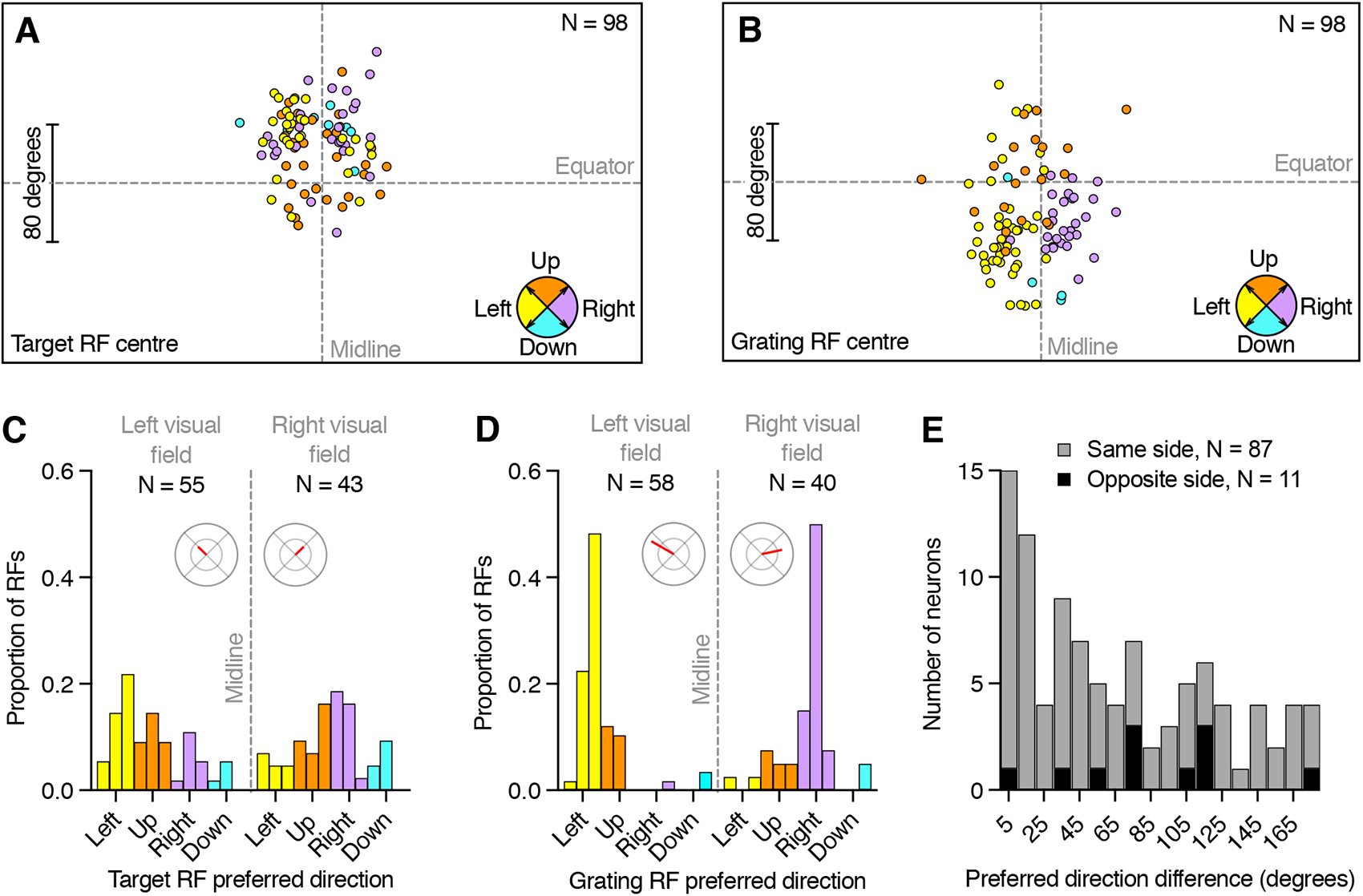
The preferred direction depends on the receptive field center location. **A)** The target receptive field centers, color coded according to their preferred direction (pictogram bottom right). **B)** The grating receptive field centers, color coded according to their preferred direction. **C)** Preferred direction of target receptive fields with centers in either the left or the right visual field (30° bins). The distribution for target receptive fields in the left visual field was significantly non-uniform (p < 0.001, Rayleigh test) with median preferred direction up and to the left, and a median vector length of 0.37 (polar plot, scale 0 to 1). The distribution for target receptive fields in the left visual field was significantly non-uniform (p = 0.0025, Rayleigh test) with median preferred direction up and to the right, and a median vector length of 0.37. **D)** Preferred direction of grating receptive fields centers in either the left or the right visual field (30° bins). The distribution for grating receptive fields in the left visual field was significantly non-uniform (p < 0.0001, Rayleigh test) with median preferred direction slightly up and to the left, and a median vector length of 0.83. The distribution for grating receptive fields in the right visual field was significantly non-uniform (p < 0.0001, Rayleigh test) with median preferred direction slightly up and to the right, and a median vector length of 0.66. **E)** Preferred direction difference between the target and grating receptive field of each neuron (10° bins). Grey data show neurons where the two receptive field centers were on the same side of the visual midline (N = 87), and black data show neurons with receptive fields on opposite sides of the visual midline (N = 11). These were not significantly different (p = 0.15, Mann-Whitney test).

The directionality of the grating receptive fields depended more strongly on center location (Fig. 3B, D). Indeed, we found a significantly non-uniform distribution (P < 0.0001, Rayleigh test), with median direction preferences slightly up and away from the visual midline (vector lengths 0.82 and 0.66, inset, Fig. 3D).

We next quantified the difference between the preferred directions of the target and the grating receptive fields. In the example neuron, this is 8° (bottom right pictogram, Fig. 2A, see also Fig. S1G, H). We found that 87 neurons had receptive field centers on the same side of the visual midline (grey, Fig. 3E), whereas 11 had receptive field centers on opposite sides (black, Fig. 3E). Across neurons the median direction difference was 75° if they were on opposite sides of the visual midline, and 44° if they were on the same side, albeit with neurons encompassing the entire span of possible directionality differences (Fig. 3E, see also Fig. S2). However, there was no distribution difference based on whether the two receptive field centers were on the same side (grey, Fig. 3E) or opposite sides (black, Fig. 3E) of the visual midline (Mann-Whitney test, P = 0.15).

### Leading-edge sensitivity matches grating receptive field

These descending neurons thus have two receptive fields, one that responds to small target motion (blue, Fig. 2, S1, S2) and one that responds to local sinusoidal gratings (red, Fig. 2, S1, S2). Which one of these receptive fields is most likely to contribute to their looming sensitivity (right, Fig. 1A-D)? Looming sensitive neurons in *Drosophila,* including the giant fiber, also respond strongly to high-contrast bars and edges (Ache et al., 2019a). We thus used full screen OFF edges to map the looming receptive field (Fig. S3). An example neuron shows strong responses to an OFF edge sweeping either left (Fig. S3A), down (Fig. S3C), or up (Fig. S3D), across the visual field, but not right (Fig. S3B). The resulting leading-edge receptive field (cyan, Fig. S3E, F) overlaps substantially with the grating receptive field (red, Fig. S3F), but not the target receptive field (blue, Fig. S3F).

Across neurons we compared the location of the leading-edge, the target, and the grating receptive field centers (circles, Fig. 4A, S3F). A qualitative analysis shows that the leading-edge receptive field centers tend to cluster below the visual equator, just like the grating receptive field centers do (cyan and red, Fig. 4A). For quantification, we calculated a proximity index (Fig. 4B). When the leading-edge receptive field center is closer to the grating receptive field center the proximity index is positive, up to a maximum of 100%. In the example neuron, the proximity index is 83% (Fig. 4B, S3). Across neurons, we found that the leading-edge receptive field centers were more frequently closer to the grating receptive field centers (11 vs 4 neurons, Fig. 4C). In those neurons where the leading-edge receptive field center was closer to the target receptive field center, the proximity index was lower (medians of -30% and 48%, Fig. 4C) and the difference was significant (Mann-Whitney test, P = 0.0015). In summary, it is likely that these neurons get their looming sensitivity predominantly within the ventral visual field, overlapping with the location of the grating receptive field.

**Figure 4.**
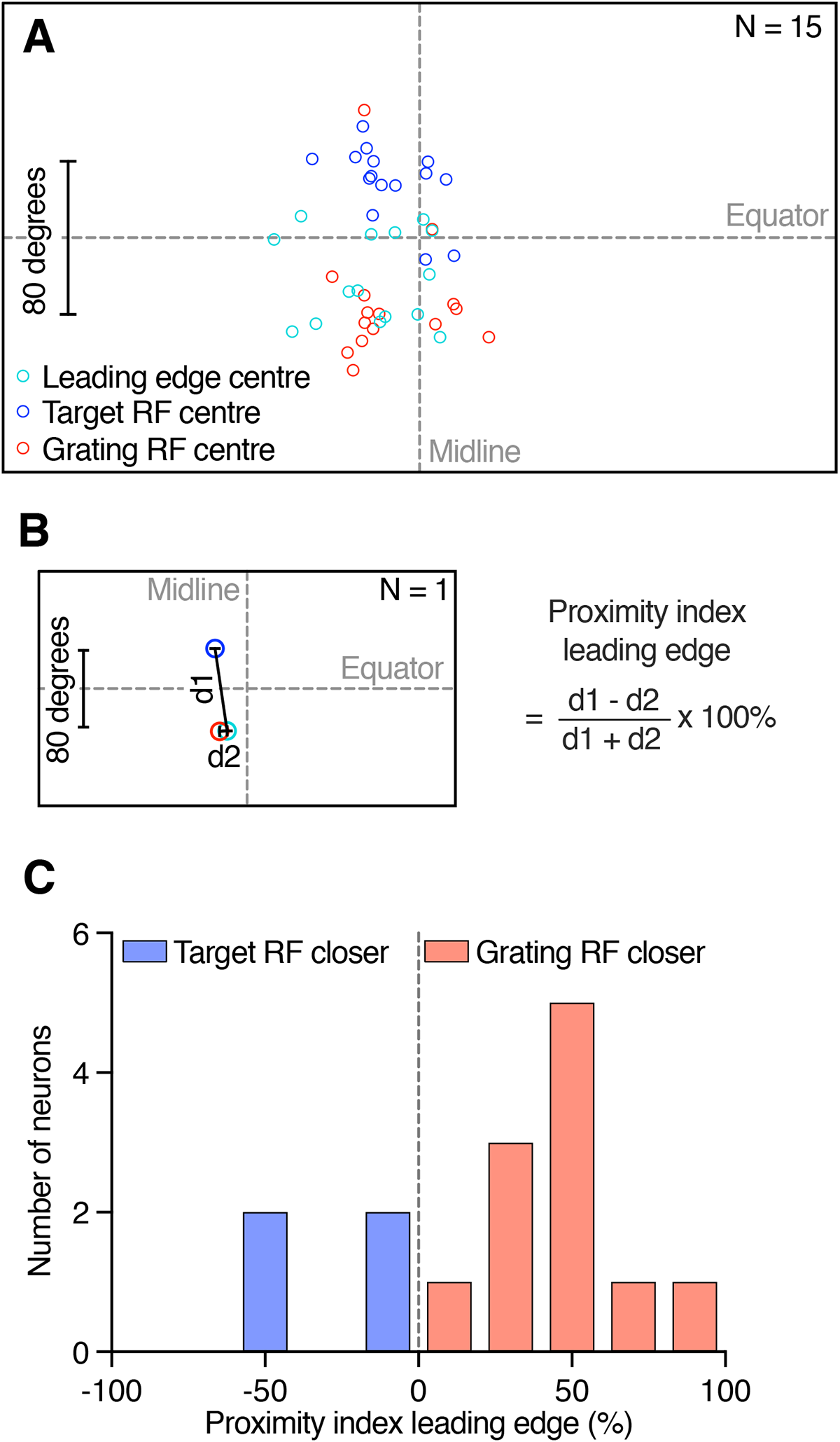
The leading-edge receptive field is closer to the grating receptive field. **A)** The location of the target (blue), grating (red) and leading-edge (cyan) receptive field centers in 15 neurons. **B)** The receptive field centers for one example neuron (*Left*), with distances (black lines) between the leading-edge and the target receptive field center (*d1*), or the grating receptive field center (*d2*), used to calculate the proximity index (*Right*). **C)** Leading edge proximity index across neurons (N = 15). The leading-edge receptive field was closer to the grating receptive field cent (red) in more neurons (N = 11) than to the target receptive field (blue, N = 4). The distribution was significantly different from 0 (P < 0.01, one sample t and Wilcoxon signed rank test), and the two distributions were different from each other (P = 0.0015, Mann-Whitney test)

### Different size response function in the two receptive fields

Previous work showed that looming sensitive neurons have a peculiar size response function, with one peak to bars of a few degrees height, similar to the size tuning of target selective neurons, and a second peak to full-screen bars (Nicholas et al., 2020). To investigate if this size sensitivity differs between the two receptive fields we scanned bars of fixed width across the visual monitor, while varying the height. We used two different trajectories, one centered on the target receptive field (blue, Fig. 5A, S4), and one on the grating receptive field (red, Fig. 5A, S4). We found that the neurons responded strongly to small bars moving through the target receptive field (blue, Fig. 5A, Fig. S4A-E), but not through the grating receptive field (red, Fig. 5A, Fig. S4A-E). For the middle-sized bars, the neurons responded weakly whether they traversed the target or the grating receptive field (Fig. 5A, S4F-I).

**Figure 5.**
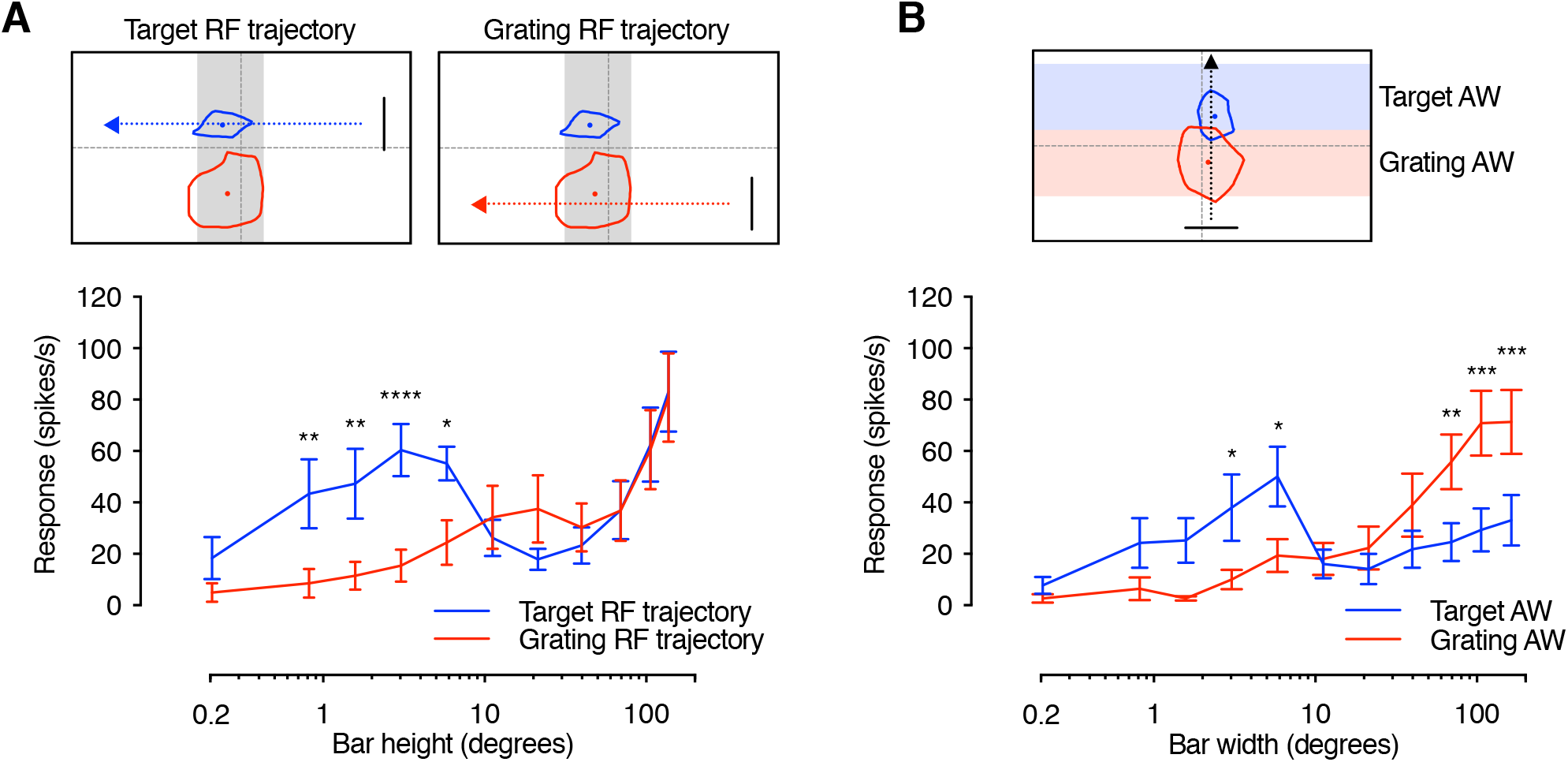
The two receptive fields have different size response functions. **A)** The pictograms indicate the bar trajectory as it moved horizontally across the screen, subtending either the target receptive field (*Left*, blue dashed line and arrow) or the grating receptive field (*Right*, red dashed line and arrow, example bar height is 84°). Typical target and grating receptive fields for an example neuron are shown. The grey shading shows the analysis window used to calculate the mean response rate, which is the same for both trajectories. The graph shows that responses to small bars are significantly stronger when passing through the target receptive field (blue) compared to the grating receptive field (red, mean ± SEM, N = 8). **B)** The pictogram indicates the analysis windows used to calculate the response to a bar of varying width as it moved vertically along the screen (trajectory in black) subtending the target receptive field (blue) or the grating receptive field (red). The graph shows that responses to narrow bars are significantly stronger within the target analysis window (AW, blue), while responses to wider bars are significantly stronger within the grating analysis window (red, mean ± SEM, N = 10). Statistical test was a two-way ANOVA followed by Sidak’s multiple comparisons, with ****P < 0.0001, ***P < 0.001, **P < 0.01 and *P < 0.05.

When the bars were extended to cover a large part of the visual monitor, they traversed both receptive fields (Fig. S4J, K), making it hard to determine which receptive field the strong response came from (Fig. 5A). To bypass this, we scanned the bars vertically instead of horizontally, so that they traversed the grating receptive field and the target receptive field at different points in time (pictogram, Fig. 5B, S5). We found that the neurons responded strongly when small bars moved through the target analysis window (blue, Fig. 5B, S5A-E), and strongly to large bars when they moved through the grating analysis window (red, Fig. 5B, S5G-L). This shows that the target receptive field is tuned to small targets, whereas the grating receptive field responds better to full-screen bars.

### Separate inputs to the two receptive fields

The data above show that looming neurons have two receptive fields (Fig. 2-4), with size tuning suggesting that they receive separate input (Fig. 5). What happens when they are stimulated simultaneously? To investigate this, we first determined the response when the two receptive fields were stimulated separately, by scanning a small target through the target receptive field (blue, Fig. 6A), and a series of bars through the grating receptive field (red, Fig. 6A, consistent with the data in Fig. 5). We compared this to the response to simultaneous stimulation (black, Fig. 6B). We found that the response to simultaneous presentation (black, Fig. 6B) was smaller than the linear sum of the two independent presentations (purple, Fig. 6B). Importantly, this cannot be due to response saturation, as the linear sum to the smallest bars (purple, Fig. 6B) is on par with the measured response to the largest bars (black, Fig. 6B).

**Figure 6.**
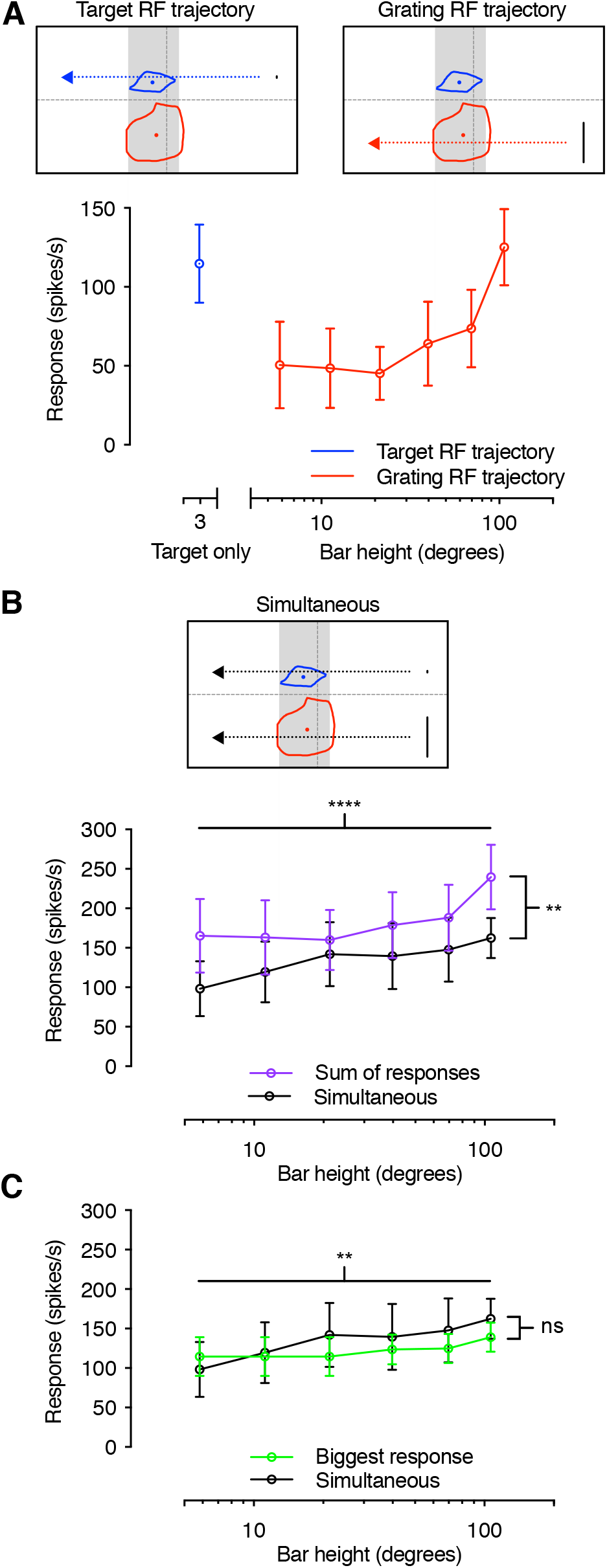
Response to simultaneous stimulation of the two receptive fields. **A)** Responses to a small target traversing the target receptive field (blue), or bars of varying heights traversing the grating receptive field (red). The pictograms at the top show the screen position of each trajectory in relation to the receptive fields for an example neuron. Grey shading indicated the analysis window used to calculate the mean response rate. **B)** Pictogram showing the screen position of simultaneously presented target and bar traversing the target and grating receptive fields. The graph shows that the responses to simultaneous presentation (black) are significantly lower than the sum of the responses to the same stimuli presented on their own (purple). **C)** The responses to simultaneous stimuli (black) are not significantly different from the strongest response evoked by either the target or the bar on its own (green). For all panels the data show mean ± SEM, for the same N = 5. Statistical analysis was done using two-way ANOVA, with ****P < 0.0001, **P < 0.01 and ns indicating P > 0.05.

We next compared the response to simultaneous presentation (black, Fig. 6C) with the strongest response for each neuron (green, Fig. 6C). We found that while there was a significant dependence on bar size, there was no significant difference between the two conditions (compare green and black, Fig. 6C, 2-way ANOVA), suggesting non-linear interactions. Similarly, the locust LGMD (Krapp and Gabbiani, 2005) displays non-linear interactions when stimuli are placed in different parts of the visual field, suggesting that the details of its receptive fields may be worth investigating in the future.

### Target receptive field is based on 1-point correlator input, whereas the grating receptive field uses 2-point correlator input

The data above show two independent receptive fields. What is the likely pre-synaptic input to each? Both optic lobe and descending target tuned neurons (Wiederman et al., 2013; Nicholas and Nordström, 2021) generate their target selectivity using 1-point correlators, which are based on the comparison of an OFF contrast change immediately followed by an ON contrast change at a single point in space (Wiederman et al., 2008). These correlators are fundamentally different from 2-point correlators, such as Hassenstein-Reichard elementary motion detectors (Hassenstein and Reichardt, 1956), in their response to high-contrast edges. For example, a 1-point correlator will respond only weakly to either OFF or ON contrast edges, compared with complete objects, whereas 2-point correlators respond equally well to single edges and complete objects (inset, Fig. 7, data replotted from Wiederman et al., 2013).

**Figure 7.**
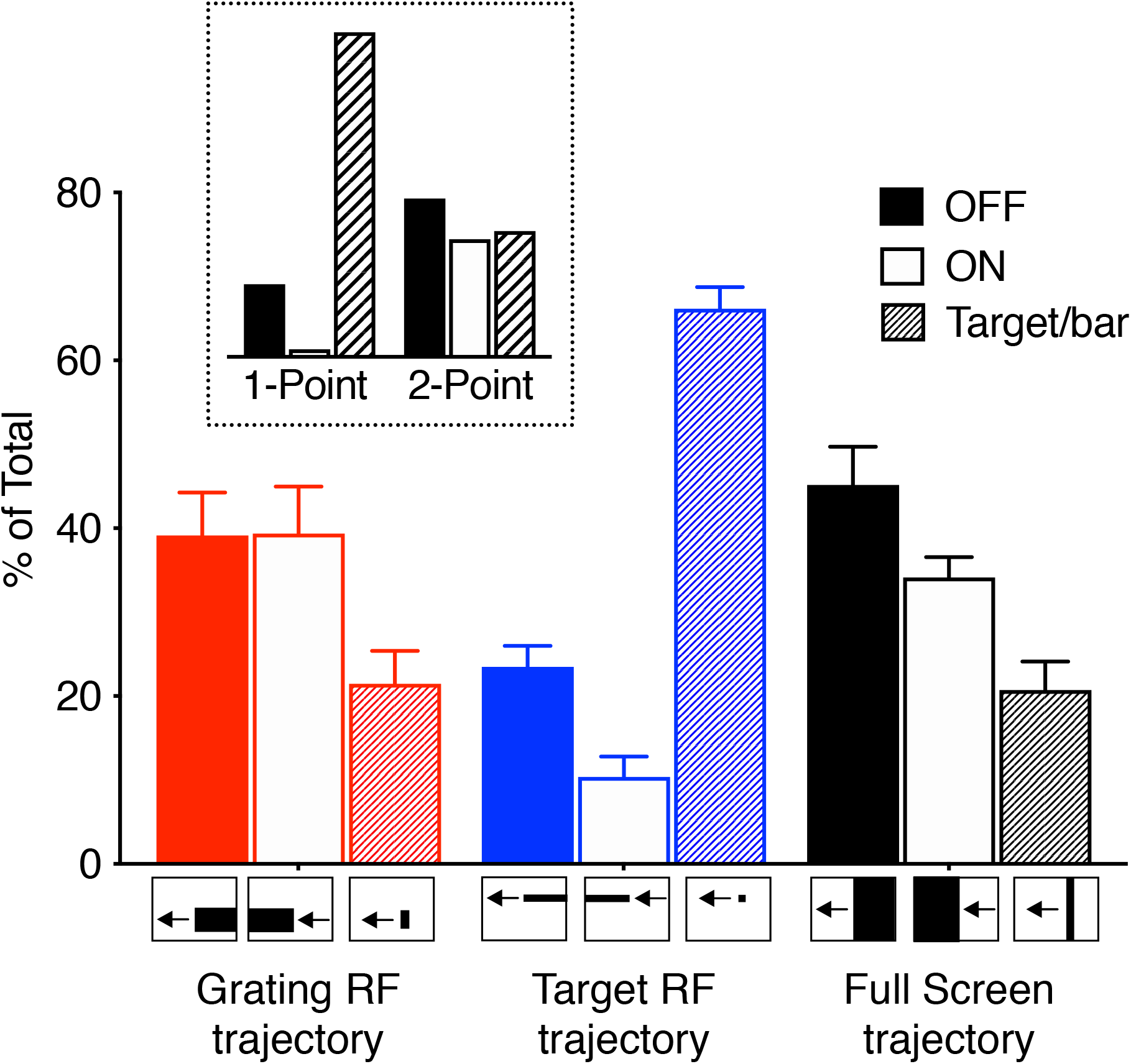
These looming neurons get input from both 1- and 2-point correlators. The response to a leading OFF edge, a trailing ON edge, or a complete bar, all with a height of 84°, traversing the grating receptive field (red, N = 9), a height of 3° traversing the target receptive field (blue, N = 10), or the full height of the screen (black, N = 7). The stimuli moved horizontally at a velocity of 209 mm/s. In all cases the response from each neuron was normalized to the sum of the response to all three stimuli from the same trajectory (i.e. OFF edge only, ON edge only, or complete bar). The inset shows the predicted response of a motion detector that compares luminance changes over one point (also referred to as an elementary STMD) or two points in space (often referred to as an EMD). The inset pictograms are replotted from Wiederman et al. (2013).

To investigate the potential input to the two receptive fields we first scanned an OFF edge through the grating receptive field, then an ON edge, followed by a complete bar (red, Fig. 7). We found that the responses to single edges were similar to the response to a complete bar (ns, red, Fig. 7). The response was thus consistent with an underlying 2-point correlator input (compare red data with inset, Fig. 7). In contrast, when we scanned edges or targets through the target receptive field, the response to a complete target was much stronger than to either OFF or ON edges on their own (one-way ANOVA, followed by Tukey’s multiple comparisons test, P < 0.0001, blue, Fig. 7). Indeed, the response was consistent with an underlying 1-point correlator input (compare blue data with inset, Fig. 7). In response to full-screen edges and bars, which cover both receptive fields, we found the strongest response to the OFF edge (P = 0.0049 for OFF vs bar, ns for ON vs bar, black, Fig. 7).

## Discussion

We have shown here a group of descending neurons in the hoverfly *Eristalis tenax* that are sensitive to both small moving targets and to looming stimuli (Fig. 1). We show that the neurons have two discrete receptive fields, with different locations (Fig. 2, S1E, F, S2) and preferred directions (Fig. 3, S1G, H, S2). We show that the looming sensitivity is likely associated with the ventral receptive field (Fig. 4, S3). The size tuning (Fig. 5, S4, S5) and sensitivity to OFF and ON contrast edges (Fig. 7) supports independent input to the two receptive fields, using two fundamentally different pre-synaptic pathways. The input from the two pathways is not linearly summed (Fig. 6B).

### Dual receptive fields

The neurons that we describe here were classified as looming sensitive based on a strong response to a looming stimulus (Fig. 1E, F) compared with an appearance control (Nicholas et al., 2020). However, as they also respond strongly to small moving targets (left, Fig. 1A-D, Fig. 5), they could have been classified as target selective descending neurons (TSDNs).

Indeed, the dragonfly TSDN DIT3 responds strongly to both small targets and to looming stimuli (Gonzalez-Bellido et al., 2013). In the locust, LGMD/DCMD neurons respond to both targets and to looming stimuli (Santer et al., 2012), and some central complex looming sensitive neurons also respond to small moving targets (Rosner and Homberg, 2013). Similarly, in *Drosophila,* some optic lobe and descending looming sensitive neurons also respond to smaller objects (e.g. de Vries and Clandinin, 2012; Klapoetke et al., 2017; Namiki et al., 2018; Ache et al., 2019a).

However, as opposed to these examples (e.g. Santer et al., 2012; Gonzalez-Bellido et al., 2013; Rosner and Homberg, 2013; Ache et al., 2019a), we show that the dual sensitivity to small targets and to larger objects is associated with two discrete receptive fields (Fig. 2 – 4, S1 – S3). It is currently unknown if the dual sensitivity described in other insects (e.g. Santer et al., 2012; Gonzalez-Bellido et al., 2013; Rosner and Homberg, 2013; Ache et al., 2019a) also comes from different receptive fields. In *Drosophila* Foma-1 target sensitivity was specific to the dorsal visual field, similar to our data (blue, Fig. 2B, C), while the visual field location of the looming sensitivity was not specified (de Vries and Clandinin, 2012).

Our recordings were done extracellularly (Fig. 1), meaning that neurons with no spontaneous activity are difficult to discover without presenting a suitable stimulus. We used a small moving target to initially identify visual neurons (left, Fig. 1A-D), thus biasing our results towards those looming sensitive descending neurons that also responded to small targets. However, it is likely that there are looming neurons that do not respond to small objects, such as found in e.g. *Drosophila* (e.g. Klapoetke et al., 2017; Ache et al., 2019b) and crabs (see e.g. Cámera et al., 2020). Additionally, our visual monitor was placed in front of the animal, thus biasing our results to neurons with frontal sensitivity. It is unlikely that the frontal clustering of receptive fields (Fig. 2B) was biased by our stimuli not being perspective distorted, as the response width did not change significantly with object size (blue data, Fig. S4). However, it is likely that there are additional looming sensitive descending neurons with dorsal receptive fields, which could be useful for e.g. detecting predators approaching from above, or lateral receptive fields, which could be useful for avoiding imminent collision. For example, in the crab there are 16 retinotopically arranged looming sensitive MLG1 neurons that underlie directional escape behaviors (Medan et al., 2015). While each receptive field is small, together the 16 neurons cover 360° of the visual field (Medan et al., 2015), and are thus able to encode directional escape responses.

A further technical limitation of our work was that we recorded from immobile animals that were placed upside down in front of the monitor. In this situation there is no feedback from the motor system, which could affect neural responses (see e.g. Fujiwara et al., 2017; Fenk et al., 2021).

### Neuronal input mechanisms

We showed that the looming sensitive descending neurons likely receive distinct input to the two receptive fields (Fig. 5-7). Indeed, the dorsal target receptive field is likely to use pre-synaptic 1-point correlators (blue, Fig. 7), just like the TSDNs do (Nicholas and Nordström, 2021). Furthermore, the size tuning of the dorsal target receptive field (blue, Fig. 5, S4-5) is similar to the size tuning of TSDNs (Nicholas et al., 2018b; Nicholas and Nordström, 2021), and of the presumably pre-synaptic STMDs (Nordström, 2012). This suggests that the dorsal target receptive field could share input with the TSDNs.

In contrast, the ventral grating receptive field responded better to larger bars than to small targets (red, Fig. 5, S4-5), similar to optic flow sensitive descending neurons (Nicholas and Nordström, 2021). In addition, the ventral grating receptive field is likely to use pre-synaptic 2-point correlators of the EMD-type (red, Fig. 7), similar to optic flow sensitive neurons (Harris et al., 1999). Interestingly, the looming sensitive LPLC2 neurons, which are pre-synaptic to the *Drosophila* giant fiber (Ache et al., 2019b), get their input from T4/T5 (Klapoetke et al., 2017), which is consistent with a 2-point, EMD-type, correlator input (see e.g. Salazar-Gatzimas et al., 2016). As our leading-edge data suggests that looming sensitivity could be associated with the grating receptive field (Fig. 4, S3), this indicates that looming sensitivity might be generated by 2-point correlation. Indeed, in the housefly, escapes can be triggered by widefield gratings, even if not as efficiently as by looming stimuli (Holmqvist and Srinivasan, 1991).

We found that the directionality of the grating receptive field depended strongly on the azimuthal location of the receptive field center (Fig. 3B, D). However, the directionality of the target receptive field was less dependent on its visual field location (Fig. 3A, C). In addition, we found that the direction preference differences of the two receptive fields covered the full 180° of possible direction differences (Fig. 3E), further supporting independent inputs.

### Behavioral role

Previous work has shown that the same stimulus displayed in different parts of the visual field can elicit different behavioral output. For example, when crabs living in mudflats see a small dummy moved at ground level they initiate prey pursuit behavior, but when the same dummy is moved above the crab, they try to escape it (Tomsic et al., 2017). In flying *Drosophila*, a looming stimulus in the lateral visual field leads to an escape response, whereas a looming stimulus in the frontal visual field leads to landing attempts (Tammero and Dickinson, 2002). While we did not stimulate the lateral visual field in our set-ups, the strong responses to frontal looming stimuli (right, Fig. 1A-D), likely associated with the ventral receptive field (Fig. 4), suggests that this could be used during landing behaviors on e.g. flowers. Indeed, bees adjust their body angle when landing so the landing surface ends up in the ventral visual field (Evangelista et al., 2010).

Alternatively, the neurons that we described here could potentially be used in pursuit. Indeed, when a hoverfly is pursuing a target, during most of the pursuit it will be projected as a small object on the pursuer’s eye (Thyselius et al., 2023). When the hoverfly is below the target, having a dorsal target receptive field would be appropriate (blue, Fig. 2A-C). This could thus be supported by either the neurons described here (blue, Fig. 2), or by TSDNs without looming sensitivity (Nicholas et al., 2018b; Nicholas et al., 2020; Nicholas and Nordström, 2021).

During later stages of the pursuit, when the hoverfly gets closer to the target (Thyselius et al., 2023), this will be seen as a looming object. It has been suggested that this part of the pursuit cannot be subserved by classic target tuned neurons, but instead requires neurons that respond to larger objects and looming stimuli (see e.g. Discussion in Bagheri et al., 2015), like in zebrafish larvae (Henriques et al., 2019). Furthermore, during the final stages before capture, the pursuer would need to orient itself to grab the target with its legs. During this stage the target would be seen as a larger object in the ventral visual field, which would make the more ventral receptive field useful (red, Fig. 2).

However, the behavioral output required during initial target detection and final capture, for predator avoidance and landing, are all quite different. The descending neurons play an important role in sensorimotor transformation (Namiki et al., 2018), but it is difficult to see how the same descending neuron could control such different behaviors. However, as behavioral state has a strong effect on visual neurons (see e.g. Fujiwara et al., 2017; Fenk et al., 2021), this could help modulate the behavioral output. For example, *Drosophila* Foma-1 controls escape responses in stationary flies (de Vries and Clandinin, 2012), as well as courtship in moving flies (Coen et al., 2016). Future work investigating the impact of internal state, as well as where the neurons described here project to, and which behaviors they thus could contribute to, will help elucidate this.

## Acknowledgements

We thank past and current lab members for constructive feedback, and the Botanic Gardens of Adelaide for their ongoing support. This research was funded by the US Air Force Office of Scientific Research (AFOSR, FA9550-19-1-0294), the Australian Research Council (ARC, DP180100144, DP210100740, DP230100006 and FT180100289).

## References

Ache JM, Namiki S, Lee A, Branson K, Card GM (2019a) State-dependent decoupling of sensory and motor circuits underlies behavioral flexibility in *Drosophila*. Nat Neurosci 22:1132–1139.

Ache JM, Polsky J, Alghailani S, Parekh R, Breads P, Peek MY, Bock DD, von Reyn CR, Card GM (2019b) Neural basis for looming size and velocity encoding in the *Drosophila* Giant Fiber escape pathway. Curr Biol 29:1073–1081.e1074.

Bagheri ZM, Wiederman SD, Cazzolato BS, Grainger S, O’Carroll DC (2015) Properties of neuronal facilitation that improve target tracking in natural pursuit simulations. J Roy Soc Interface 12:20150083.

Berens P (2009) CircStat: A MATLAB toolbox for circular statistics. Journal of Statistical Software 31:1–21.

Brainard DH (1997) The Psychophysics toolbox. Spatial Vision 10:433–436.

Cámera A, Belluscio MA, Tomsic D (2020) Multielectrode recordings from identified neurons involved in visually elicited escape behavior. Front Behav Neurosci 14:592309.

Coen P, Xie M, Clemens J, Murthy M (2016) Sensorimotor transformations underlying variability in song intensity during *Drosophila* courtship. Neuron 89:629–644.

de Vries SE, Clandinin TR (2012) Loom-sensitive neurons link computation to action in the *Drosophila* visual system. Curr Biol 22:353–362.

Evangelista C, Kraft P, Dacke M, Reinhard J, Srinivasan MV (2010) The moment before touchdown: landing manoeuvres of the honeybee *Apis mellifera*. J Exp Biol 213:262–270.

Fabian ST, Sumner ME, Wardill TJ, Rossoni S, Gonzalez-Bellido PT (2018) Interception by two predatory fly species is explained by a proportional navigation feedback controller. J Roy Soc Interface 15:20180466.

Fenk LM, Kim AJ, Maimon G (2021) Suppression of motion vision during course-changing but not course-stabilizing navigational turns. Curr Biol 31:608–4619.

Fitzpatrick SM, Wellington WG (1983) Contrasts in the territorial behavior of three species of hover flies (Diptera: Syrphidae). Can Entomol 115:559–566.

Fotowat H, Gabbiani F (2007) Relationship between the phases of sensory and motor activity during a looming-evoked multistage escape behavior. J Neurosci 27:10047–10059.

Fotowat H, Fayyazuddin A, Bellen HJ, Gabbiani F (2009) A novel neuronal pathway for visually guided escape in *Drosophila melanogaster*. J Neurophysiol 102:875–885.

Frye MA, Olberg RM (1995) Visual receptive field properties of feature detecting neurons in the dragonfly. J Comp Physiol A 177:569–576.

Fujiwara T, Cruz TL, Bohnslav JP, Chiappe ME (2017) A faithful internal representation of walking movements in the *Drosophila* visual system. Nat Neurosci 20:72–81.

Gonzalez-Bellido PT, Peng H, Yang J, Georgopoulos AP, Olberg RM (2013) Eight pairs of descending neurons in the dragonfly give wing motor centers accurate population vector of prey direction. Proc Natl Acad Sci USA 110:676–701.

Harris RA, O’Carroll DC, Laughlin SB (1999) Adaptation and the temporal delay filter of fly motion detectors. Vision Res 39:2603–2613.

Hassenstein B, Reichardt W (1956) Systemtheoretische Analyse der Zeit, Reihenfolgen und Vorzeichenauswertung Bei der Bewegungsperzeption des Rüsselkafers Chlorophanus. Z Naturforsch 11: 513–524.

Henriques PM, Rahman N, Jackson SE, Bianco IH (2019) Nucleus isthmi is required to sustain target pursuit during visually guided prey-catching. Curr Biol 29:1771–1786.e1775.

Holmqvist MH, Srinivasan MV (1991) A visually evoked escape response of the housefly. J Comp Physiol A 169:451–459.

Klapoetke NC, Nern A, Peek MY, Rogers EM, Breads P, Rubin GM, Reiser MB, Card GM (2017) Ultra-selective looming detection from radial motion opponency. Nature 551:237–241.

Krapp HG, Gabbiani F (2005) Spatial distribution of inputs and local receptive field properties of a wide-field, looming sensitive neuron. J Neurophysiol 93:2240–2253.

Leibbrandt R, Nicholas S, Nordström K (2021) The impulse response of optic flow-sensitive descending neurons to roll m-sequences. J Exp Biol 224.

Lenzi SC, Cossell L, Grainger B, Olesen SF, Branco T, Margrie TW (2022) Threat history controls flexible escape behavior in mice. Curr Biol 32:2972–2979.e2973.

Liu YJ, Wang Q, Li B (2011) Neuronal responses to looming objects in the superior colliculus of the cat. Brain Behav Evol 77:193–205.

Mancienne T, Marquez-Legorreta E, Wilde M, Piber M, Favre-Bulle I, Vanwalleghem G, Scott EK (2021) Contributions of luminance and motion to visual escape and habituation in larval zebrafish. Front Neural Circuits 15:748535.

Medan V, Beron De Astrada M, Scarano F, Tomsic D (2015) A network of visual motion-sensitive neurons for computing object position in an arthropod. J Neurosci 35:6654–6666.

Nakagawa H, Hongjian K (2010) Collision-sensitive neurons in the optic tectum of the bullfrog, *Rana catesbeiana*. J Neurophysiol 104:2487–2499.

Namiki S, Dickinson MH, Wong AM, Korff W, Card GM (2018) The functional organization of descending sensory-motor pathways in *Drosophila*. eLife 7:e34272.

Nicholas S, Nordström K (2021) Facilitation of neural responses to targets moving against optic flow. Proc Natl Acad Sci U S A 118.

Nicholas S, Leibbrandt R, Nordström K (2020) Visual motion sensitivity in descending neurons in the hoverfly. J Comp Physiol A 206:149–163.

Nicholas S, Thyselius M, Holden M, Nordström K (2018a) Rearing and long-term maintenance of *Eristalis tenax* hoverflies for research studies. JoVE:e57711.

Nicholas S, Supple J, Leibbrandt R, Gonzalez-Bellido PT, Nordström K (2018b) Integration of small- and wide-field visual features in Target-Selective Descending Neurons of both predatory and non-predatory dipterans. J Neurosci 38:10725–10733.

Nityananda V, Tarawneh G, Rosner R, Nicolas J, Crichton S, Read J (2016) Insect stereopsis demonstrated using a 3D insect cinema. Sci Rep 6:18718.

Nordström K (2012) Neural specializations for small target detection in insects. Curr Opin Neurobiol 22:272–278.

Nordström K, O’Carroll DC (2009) The motion after-effect: Local and global contributions to contrast sensitivity. Proc R Soc Lond B 276:1545–1554.

Nordström K, Barnett PD, O’Carroll DC (2006) Insect detection of small targets moving in visual clutter. PLoS Biol 4:378–386.

Olberg RM, Seaman RC, Coats MI, Henry AF (2007) Eye movements and target fixation during dragonfly prey-interception flights. J Comp Physiol A 193:685–693.

Pelli DG (1997) The VideoToolbox software for visual psychophysics: Transforming numbers into movies. Spatial Vision 10:437–442.

Rind FC, Simmons PJ (1992) Orthopteran DCMD neuron: A re-evaluation of responses to moving objects. I. Selective responses to approaching objects. J Neurophysiol 68:1654–1666.

Rosner R, Homberg U (2013) Widespread sensitivity to looming stimuli and small moving objects in the central complex of an insect brain. J Neurosci 33:8122–8133.

Salazar-Gatzimas E, Chen J, Creamer MS, Mano O, Mandel HB, Matulis CA, Pottackal J, Clark DA (2016) Direct measurement of correlation responses in *Drosophila* elementary motion detectors reveals fast timescale tuning. Neuron 92:227–239.

Santer RD, Rind FC, Simmons PJ (2012) Predator versus prey: locust looming-detector neuron and behavioural responses to stimuli representing attacking bird predators. PLoS One 7:e50146.

Santer RD, Yamawaki Y, Rind FC, Simmons PJ (2008) Preparing for escape: an examination of the role of the DCMD neuron in locust escape jumps. J Comp Physiol A Neuroethol Sens Neural Behav Physiol 194:69–77.

Tammero LF, Dickinson MH (2002) Collision-avoidance and landing responses are mediated by separate pathways in the fruit fly, *Drosophila melanogaster*. J Exp Biol 205:2785–2798.

Temizer I, Donovan JC, Baier H, Semmelhack JL (2015) A visual pathway for looming-evoked escape in larval zebrafish. Curr Biol 25:1823–1834.

Thyselius M, Ogawa Y, Leibbrandt R, Wardill TJ, Gonzalez-Bellido PT, Nordström K (2023) Hoverfly (*Eristalis tenax*) pursuit of artificial targets. J Exp Biol 226:jeb244895.

Tomsic D, Sztarker J, Berón de Astrada M, Oliva D, Lanza E (2017) The predator and prey behaviors of crabs: from ecology to neural adaptations. J Exp Biol 220:2318–2327.

von Reyn CR, Nern A, Williamson WR, Breads P, Wu M, Namiki S, Card GM (2017) Feature integration drives probabilistic behavior in the *Drosophila* escape response. Neuron 94:1190–1204.e1196.

Wiederman SD, Shoemaker PA, O’Carroll DC (2008) A model for the detection of moving targets in visual clutter inspired by insect physiology. PLoS ONE 3:e2784.

Wiederman SD, Shoemaker PA, O’Carroll DC (2013) Correlation between OFF and ON channels underlies dark target selectivity in an Insect visual system. J Neurosci 33:13225–13232.

Zeil J (1986) The territorial flight of male houseflies (*Fannia canicularis* L.). Behav Ecol Sociobiol 19:312–219.

