## Supplemental pdf for "Dual receptive fields underlying target and wide-field motion sensitivity in looming sensitive descending neurons"

### Supplementary Information

***Figure S1. Receptive field mapping method.*** **A)** The resulting 20 x 20 analysis windows after scanning the entire monitor along 20 evenly spaced horizontal and vertical target trajectories. The blue boxes highlight the positions of the analysis windows (*i* - *iiii*) shown in panel C. **B)** The resulting 8 x 6 grid of overlapping patches of sinusoidal gratings. The red boxes highlight the grid locations (*i* - *iiii*) shown in panel D. **C)** The responses (open symbols) to 4 directions of target motion within the analysis windows highlighted in panel A. We fit a cosine function (black) to the responses and used this to extract the local preferred direction (LPD, filled triangle), and local motion sensitivity (LMS, vertical bar). The dotted line shows the mean response. **D)** The responses (open symbols) to 8 different directions within the positions highlighted in panel B, the resulting cosine function (black), the local preferred direction (LPD, filled triangle), local motion sensitivity (LMS, vertical bar), and mean response (dotted line). **E)** The target receptive field's position, shown as contour lines (blue and grey). The centre position (blue circle) was defined as the centroid of the 50% contour line. **F)** The grating receptive field's position, shown as contour lines (red and grey), and its centre (red circle). **G)** The arrows show the local motion sensitivity (arrow length) and local preferred direction (arrow angle) of the target receptive field. Blue arrows indicate the preferred direction in positions where the local motion sensitivity was greater than 50% of the maximum for this neuron. Bottom right pictogram indicates the resulting average preferred direction. **H)** Local motion sensitivity and preferred direction of the grating receptive field. The bottom right pictogram indicates the average preferred direction (red). The data in this figure comes from the example neuron shown in Figure 2A.

***Figure S2. Looming sensitive descending neurons are a heterogeneous group of neurons.*** **A)** Target (blue) and grating (red) receptive field of an example neuron. The contour lines show the 50% response, and the circles the centres of each receptive field. The arrows show the local preferred direction for positions where the response was >50% of the maximum local motion sensitivity. The bottom right pictogram indicates the average preferred direction for the target receptive field (blue) and grating receptive field (red), as well as the difference between the two (black arc and value). The Euclidean distance between the two receptive field centres (black line and value) is indicated on the left. **B)** Example of a looming sensitive neuron with small target and grating receptive fields, and a large distance between the centres. **C)** Example neuron with a large difference between their preferred directions. **D)** Example neuron where the grating receptive field is dorsal to the target receptive field. **E)** Example neuron where the grating receptive field is dorsal to the equator and overlapping with the target receptive field. **F)** Example neuron where the grating and target receptive fields are large and partially overlapping.

***Figure S3. Leading edge receptive field method.*** **A)** Example response from a single neuron (mean +/- SEM) to a full screen OFF edge moving left across the entire width of the screen. Dashed purple line indicates 50% of the maximum response achieved to any direction of leading-edge motion in this neuron. Thresholds (cyan lines) are defined as the time when the neuronal response passes 50% of the maximal response. **B)** Neuronal response to an OFF edge moving right. As the response did not reach 50% maximum, thresholds were not achieved in this direction. **C)** Neuronal response to an OFF edge moving down. **D)** The example neuron showed maximal response (solid purple line) to an OFF edge moving up. **E)** Leading edge receptive field (shaded cyan region), based on screen position of stimulus when thresholds were reached (dashed cyan lines). **F)** Screen position for grating (red), target (blue) and leading edge (cyan) receptive fields, for this example neuron. The data in panels A-D have been smoothed with a 100 ms square-wave filter and is shown at 40 kHz resolution.

***Figure S4. Response to small bars is stronger in the dorsal, target receptive field.*** Response from a single example neuron to a bar of varying height traversing the entire width of the screen (mean ± SEM, n = 4) across either the target receptive field (blue) or the grating receptive field (red). The 50% contour lines and centres for the target receptive field (blue) and grating receptive field (red) for this neuron are shown. Grey shading indicates the analysis window used to calculate the mean response for this neuron in Figure 5A. Bar heights for each panel (A) 0.2°, (B) 0.8°, (C) 1.6°, (D) 3.1°, (E) 5.7°, (F) 11.2°, (G) 21.2°, (H) 39.7°, (I) 69.7°, (J) 106.5°, (K) 137.6°. The data have been smoothed with a 100 ms square-wave filter, and is shown at 40 kHz resolution.

***Figure S5. Sensitivity to bar width depends on which receptive field it is in.*** Response from an example neuron to a bar of varying width traversing the entire height of the screen (mean ± SEM, n = 4). Pictograms indicate the position of the bar trajectory as it moved vertically up the screen (black dashed line and arrow), traversing first the grating receptive field (red) and then the target receptive field (blue). Shading indicates the two analysis windows used to calculate the mean neuronal response for this neuron in Figure 5B. Bar width for each panel (A) 0.2°, (B) 0.8°, (C) 1.6°, (D) 3.1°, (E) 5.7°, (F) 11.2°, (G) 21.2°, (H) 39.7°, (I) 69.7°, (J) 106.5°, (K) 137.6°, (L) 155.4°. The data have been smoothed with a 100 ms square-wave filter, and is shown at 40 kHz resolution.

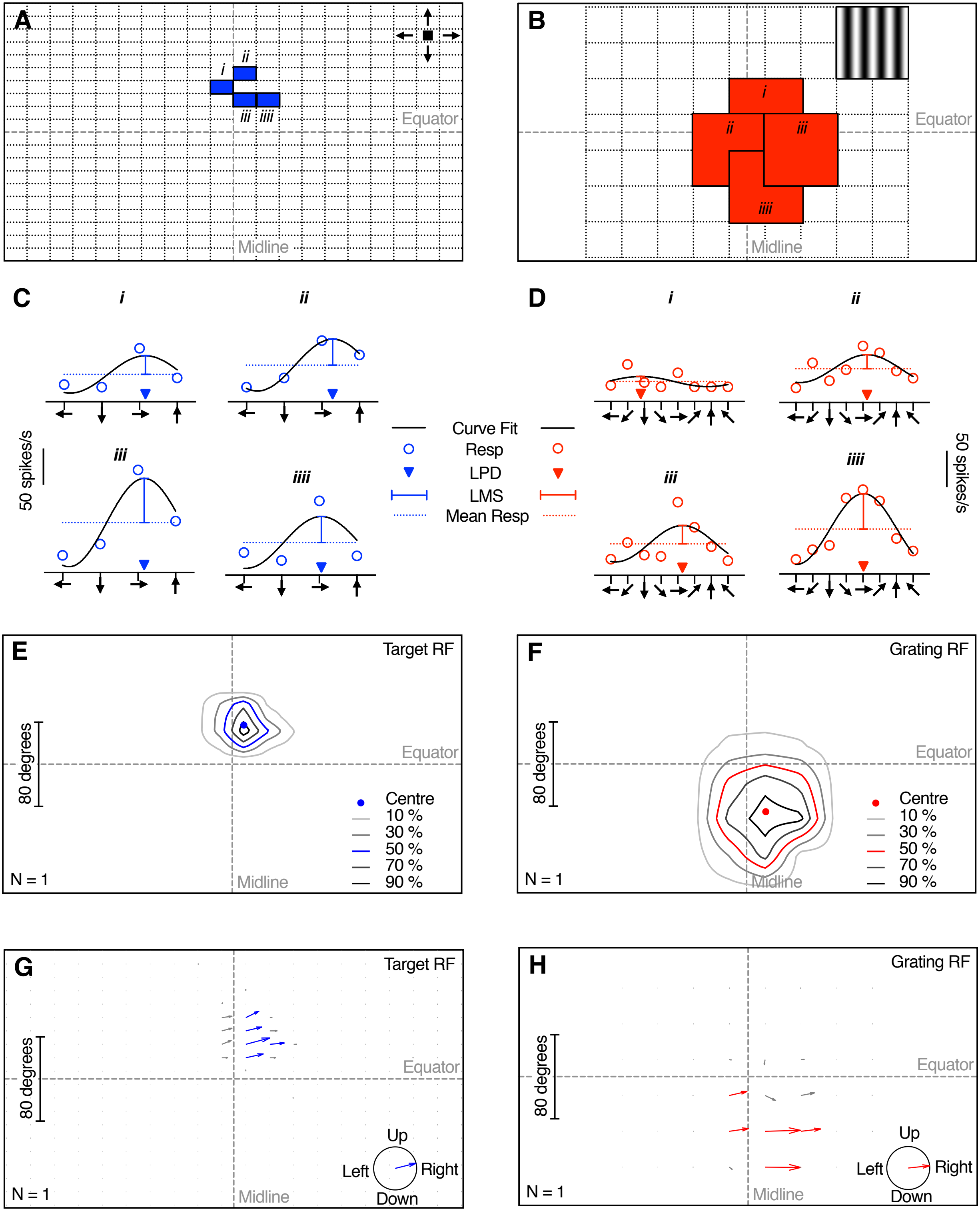

A
i
ii
iii
iiii
Equator
Midline
B
i
ii
iii
iiii
Equator
Midline
C
i
ii
iii
iiii
50 spikes/s
Curve Fit
Resp
LPD
LMS
Mean Resp
D
i
ii
iii
iiii
50 spikes/s
E
Target RF
80 degrees
Equator
Centre
10 %
30 %
50 %
70 %
90 %
N = 1
Midline
F
Grating RF
80 degrees
Equator
Centre
10 %
30 %
50 %
70 %
90 %
N = 1
Midline
G
Target RF
80 degrees
Equator
Up
Left
Right
Down
N = 1
Midline
H
Grating RF
80 degrees
Equator
Up
Left
Right
Down
N = 1
Midline

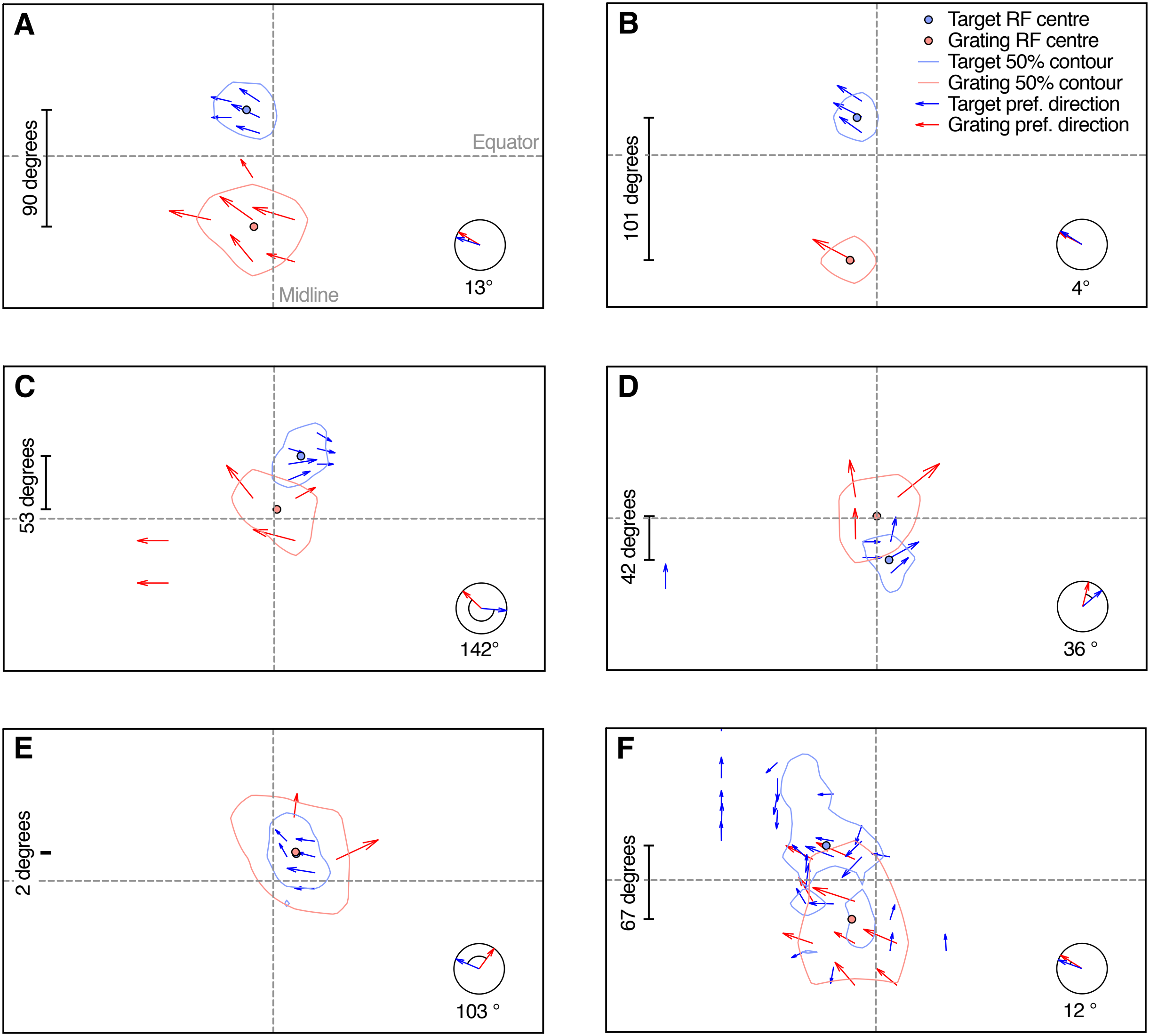

A
90 degrees
Equator
Midline
13°
B
Target RF centre
Grating RF centre
Target 50% contour
Grating 50% contour
Target pref. direction
Grating pref. direction
101 degrees
4°
C
53 degrees
142°
D
42 degrees
36 °
E
2 degrees
103 °
F
67 degrees
12 °

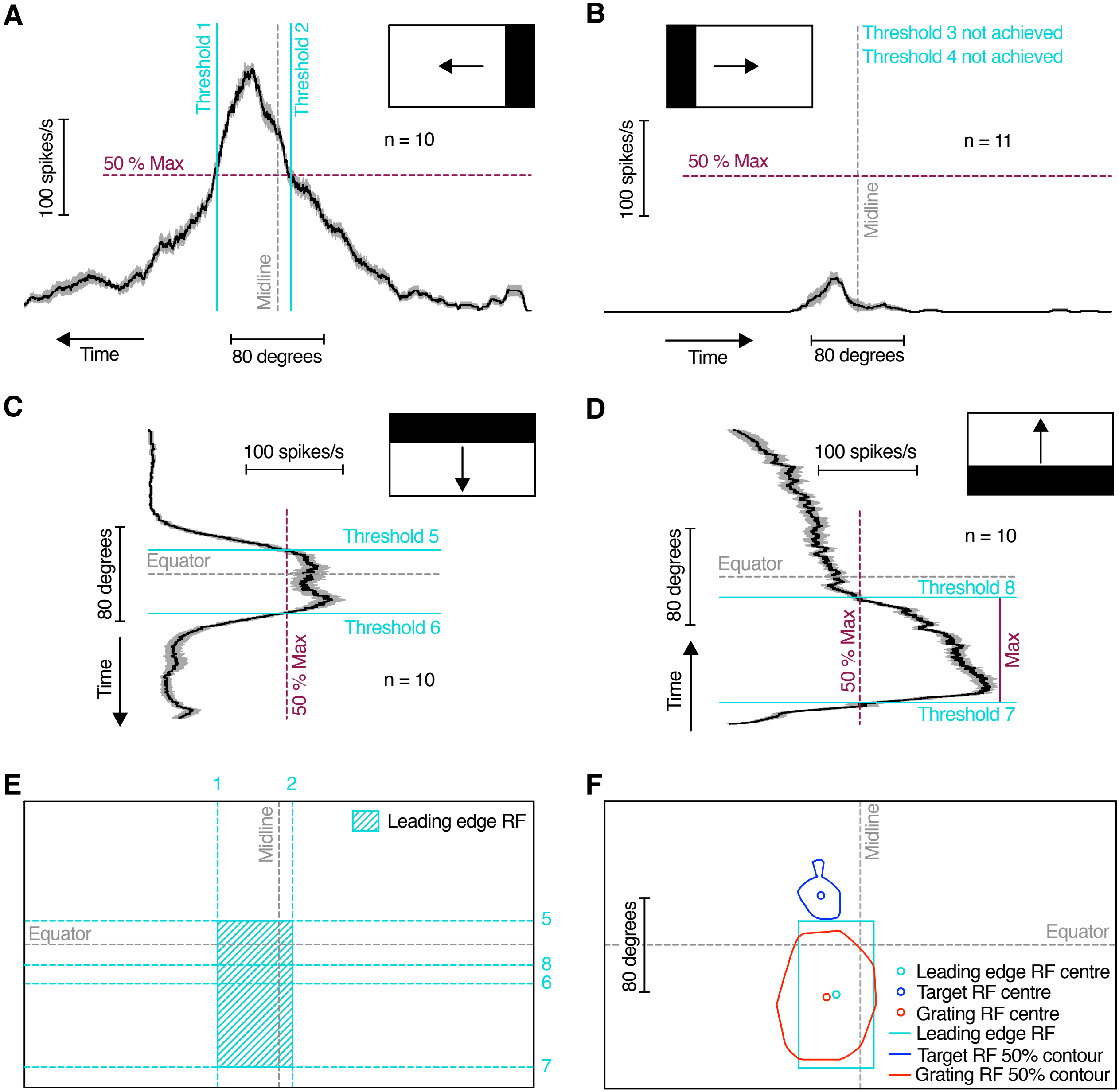

A
Threshold 1
Threshold 2
100 spikes/s
50 % Max
Midline
n = 10
Time
80 degrees
B
Threshold 3 not achieved
Threshold 4 not achieved
100 spikes/s
50 % Max
Midline
n = 11
Time
80 degrees
C
100 spikes/s
80 degrees
Equator
Threshold 5
Threshold 6
Time
50 % Max
n = 10
D
100 spikes/s
n = 10
80 degrees
Equator
Threshold 8
50 % Max
Max
Time
Threshold 7
E
1
2
Midline
Leading edge RF
Equator
5
8
6
7
F
Midline
80 degrees
Equator
Leading edge RF centre
Target RF centre
Grating RF centre
Leading edge RF
Target RF 50% contour
Grating RF 50% contour

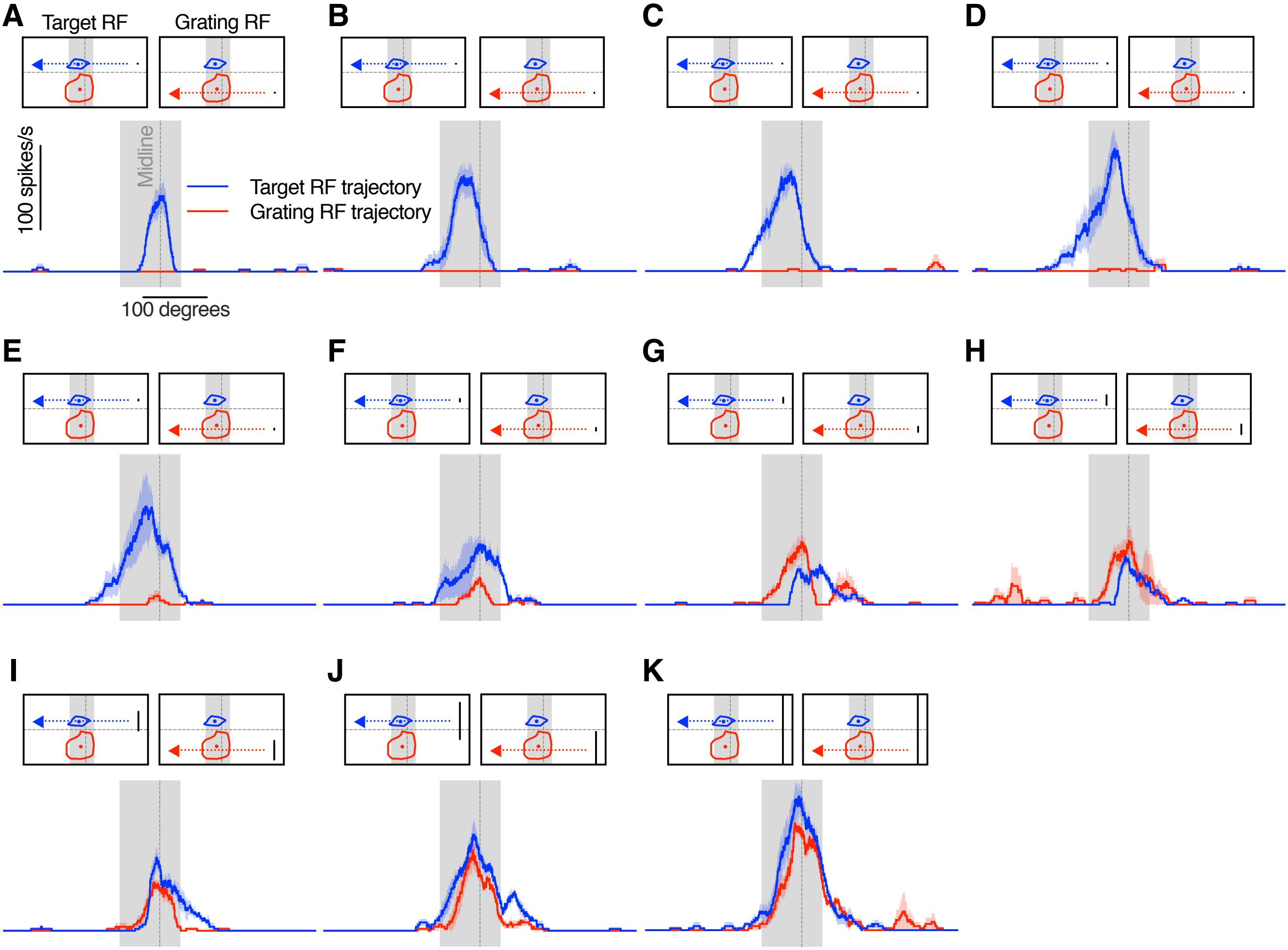
A
Target RF
Grating RF
B
C
D
100 spikes/s
Midline
Target RF trajectory
Grating RF trajectory
100 degrees
E
F
G
H
I
J
K

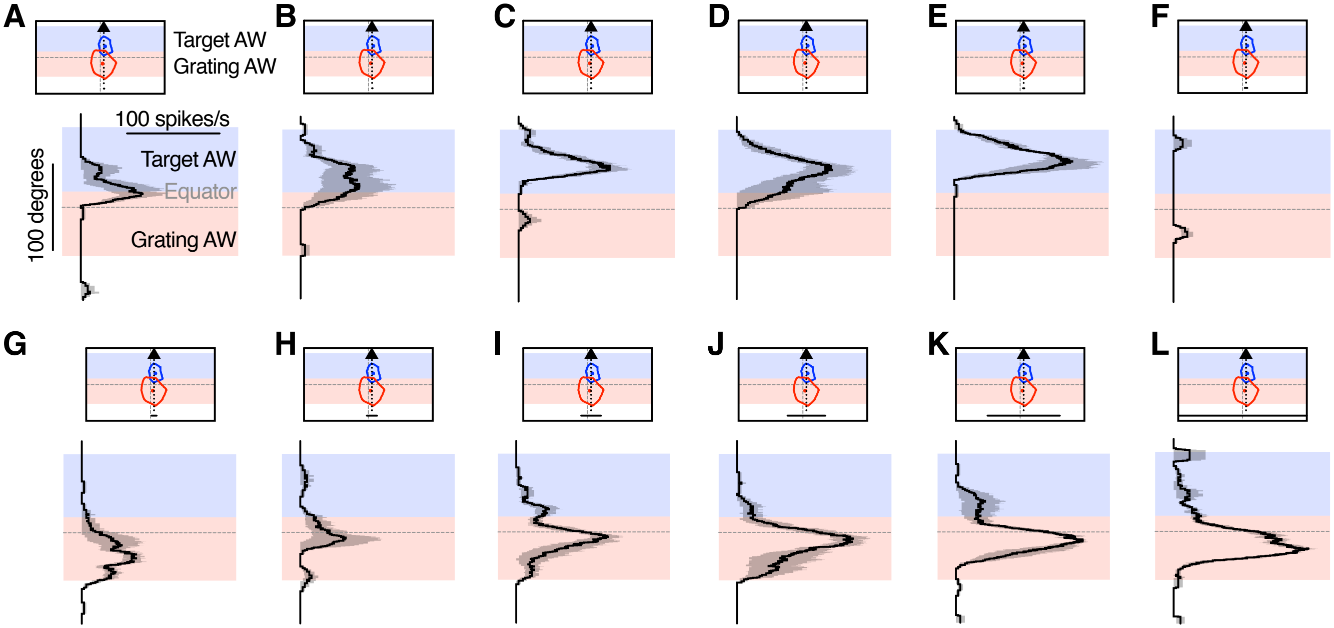
A
B
C
D
E
F
G
H
I
J
K
L
Target AW
Grating AW
100 spikes/s
Target AW
Equator
Grating AW
100 degrees
